# Buffered EGFR signaling regulated by *spitz* to *argos* expression ratio is critical for patterning the *Drosophila* eye

**DOI:** 10.1101/2021.05.19.444784

**Authors:** Nikhita Pasnuri, Manish Jaiswal, Krishanu Ray, Aprotim Mazumder

**Author notes:** <u>***Correspondence:**</u>.

## Abstract

The Epidermal Growth Factor Receptor (EGFR) signaling pathway plays a critical role in regulating tissue patterning. *Drosophila* EGFR (DER) signaling achieves specificity through multiple ligands and feedback loops to finetune signaling spatiotemporally. The principal *Drosophila* EGF, cleaved Spitz, and the negative feedback molecule, Argos are diffusible and can act both in a cell autonomous and non-autonomous manner. The relative expression dose of Spitz and Argos early in development has been shown to be critical in patterning the *Drosophila* eye, but the exact identity of the cells expressing these genes in the larval eyedisc has been elusive. Using single molecule RNA Fluorescence in situ Hybridization (smFISH), we reveal an intriguing differential expression of *spitz* and *argos* in the *Drosophila* third instar eye imaginal disc indicative of directional DER signaling. By genetically tuning DER signaling, we show that rather than absolute levels of expression, the ratio of expression to be critical for determining the adult eye phenotype. Proper ommatidial patterning is robust to thresholds around a tightly maintained wildtype ratio, and breaks down beyond. This provides a powerful instance of developmental buffering.

## Introduction

Receptor tyrosine kinases (RTKs) are key regulators of diverse cellular processes and development. Mutations or aberrant activation/inactivation of RTKs lead to different anomalies including cancers (Lemmon and Schlessinger, 2010). RTKs trigger a cytoplasmic signaling cascade involving Ras-MAPK pathway (Sundaram, 2006). EGF receptor signaling is extensively studied for its role in cellular homeostasis and in relation to human diseases (Sundaram, 2006). *Drosophila melanogaster* has one EGF receptor (DER) and four activating ligands (Shilo, 2014). The difference in expression of the ligands in a tight spatiotemporal pattern was suggested to bring about difference in EGF responses. Among *Drosophila* EGFR ligands, Gurken (Ghiglione et al., 2002; Neuman-Silberberg and Schüpbach, 1993; Schüpbach, 1987), Spitz (Mayer and Nüsslein-Volhard, 1988; Schweitzer et al., 1995a; Tio and Moses, 1997) and Keren (Reich and Shilo, 2002) are homologous to TGFα and Vein (Donaldson et al., 2004; Schnepp et al., 1996; Simcox et al., 1996) is homologous to neuregulin. Spitz is the canonical EGF in the fly. Beyond different ligands, positive and negative feedback loops are known to tightly regulate EGFR signaling (Brandman and Meyer, 2008; Perrimon et al., 2012). The downstream targets of DER include activators like Vein and Rhomboid (Bier et al., 1990; Urban et al., 2002) and inhibitors like Argos (Golembo et al., 1996; Schweitzer et al., 1995b; Wasserman and Freeman, 1998a), Kekkon-1 (Ghiglione, 2003; Ghiglione et al., 1999), Sprouty (Casci et al., 1999; Hacohen et al., 1998). Of the feedback molecules, Sprouty, Rhomboid and Kekkon-1 act in a cell autonomous manner, while Argos is a diffusible factor like cleaved Spitz, and can potentially act both in a cell autonomous and non-autonomous manner. The competitive and stoichiometric sequestration of Spitz by Argos has been shown to be critical for patterning different tissues (Schweitzer et al., 1995b; Taguchi et al., 2000). Spitz and Argos are thought to be short-range activator and long-range inhibitor respectively (Freeman, 1997).

A strong instance of EGFR-mediated patterning occurs in the *Drosophila* compound eye with its periodic units called ommatidia. A single ommatidium comprises of eight neuronal photoreceptors (PR) accompanied by twelve non-neuronal cells (Baker and Yu, 2001; Kumar, 2012; Waddington, C.H. and Perry, 1969). The photoreceptor differentiation starts in the early 3^rd^ instar larvae along an anteriorly progressing wave of Hedgehog in the eye-antennal imaginal disc leaving a morphogenetic furrow behind (Greenwood and Struhl, 1999). Posterior PR cells are older in developmental time than clusters just behind the morphogenetic furrow. EGFR signaling is a prerequisite for photoreceptor differentiation and also cone and pigment cells until pupal stage (Freeman, 1996; Tio and Moses, 1997).

Spitz binds to DER and activates the cytoplasmic Ras-MAPK pathway (Ambrosio et al., 1989; Brand and Perrimon, 1994). This cascade activates the transcriptional activator, PntP1 and degrades transcriptional repressor, Yan (Lai and Rubin, 1992; O’Neill et al., 1994; Rebay and Rubin, 1995). Downstream of the signaling pathway, *argos* is expressed and the protein product is secreted out of the cell. Argos binds Spitz in a 1:1 ratio by clamping to the EGF domain and restricts the amount of free ligand available for DER activation (Klein et al., 2004; Klein et al., 2008). Dosage of EGFR components is known to maintain a biochemical balance, which dictates the final strength of the signaling pathway. Halving the dosage of Spitz and Argos gave rise to a patterned ommatidial arrangement whereas halving only Argos or overexpressing Argos gave rise to a rough eye phenotype (Schweitzer et al., 1995b; Taguchi et al., 2000). The relative strength of signaling decides the cellular choice, which in turn contributes towards proper pattern formation. But while relative expression dose of Spitz and Argos early in development has been shown to be critical in patterning the *Drosophila* eye, the exact identity of the cells expressing these genes in the larval eyedisc has been elusive.

Argos is both a negative regulator and a target of EGFR signaling. *argos* expression is a specific proxy for strong EGFR signaling (Gabay, 1997; Wasserman and Freeman, 1998b). To quantitatively understand the level of expression of diffusible DER components, and the dosage of EGFR signaling following the protein products can be problematic. Immunofluorescence staining or reporter trap lines to track diffusible proteins is challenging as the inference on the type of cells which have secreted them is not possible (Gabay, 1997). The dual phosphorylated ERK (dpERK) staining is classically used to read the strength of the EGFR pathway. Since most of the RTK pathways converge on the MAPK pathway, dpERK staining may not specifically and quantitatively report on EGFR strength. These problems can be circumvented by detecting endogenous mRNA *in situ*, which additionally could also indicate the identity of cells that express these genes. In recent years Single-molecule Fluorescence *In Situ* Hybridization (smFISH) has emerged as a powerful method to sensitively and quantitatively report the gene expression in wholemount tissues, along with cell-to-cell variability (Pasnuri et al., 2018; Trcek et al., 2017; Yang et al., 2017). We have previously shown that such methods can be used to detect even low levels of gene expression in the wholemount *Drosophila* tissues (Pasnuri et al., 2018).

In this paper, we show the directional EGFR signaling from the photoreceptors to the neighboring uncommitted cells using smFISH for EGFR pathway genes. We reveal an intriguing differential expression of *spitz* and *argos* in photoreceptor and non-photoreceptor cells of the larval eye disc. We show that relative expression levels of *spitz* and *argos* is important for the generation of proper ommatidial pattern rather than their absolute expression level. By systematically tuning the expression of EGFR pathway genes, we analyze the biochemical buffer range where the relative gene expression levels of the ligand *spitz* and the negative feedback molecule *argos* contribute towards proper pattern formation in the *Drosophila* eye.

## Results

### Directional EGFR signaling in the eye imaginal disc

EGFR and the principal EGF, Spitz in *Drosophila* are expressed uniformly throughout multiple tissues during the development (Zak and Shilo, 1992). In the 3^rd^ instar eye imaginal disc, Spitz is responsible for activation of EGFR signaling posterior to the morphogenetic furrow. Downstream targets of EGFR signaling like *argos* are expressed as a downstream response to PntP1 activation (Shwartz et al., 2013). As Spitz and Argos are diffusible factors that can act in both a cell autonomous and non-autonomous manner and act in a 1:1 stoichiometry to modulate EGFR signaling (Klein et al., 2004), we investigated their transcription status in cells behind the morphogenetic furrow using smFISH. We used Elav, a pan-neuronal marker (Robinow and White, 1991) to distinguish photoreceptor (PR) cells from other non-photoreceptor and undifferentiated pool of cells. In-situ hybridization (ISH) for *spitz* has been performed in eye discs at this stage (Tio and Moses, 1997), and dpERK patterns also described (Gabay, 1997). Despite this, clear cell type-specific differences have not been highlighted most likely due to lower sensitivity of the methods used. Using smFISH, somewhat to our surprise we found that *spitz* is clearly expressed in higher levels in the photoreceptor cells (Elav-positive cells) compared to neighboring cells (Fig. 1A, 1B and Supp. Fig. 1B). Our methods allow us to follow two different mRNA species in the same tissue, and we found that the downstream target gene, *argos*, is expressed exclusively in the neighboring cells which have not yet made a cell fate choice (Fig. 1A, 1B and Supp. Fig. 1B). Because *argos* expression is direct target and thus a proxy for strong EGFR signaling, this indicates the exclusively cell non-autonomous effect of Spitz secreted by photoreceptor cells on their neighboring cells. Since the nuclei are densely packed in the eye imaginal disc, we quantified the mRNA expression and normalized to a 1000μm^3^ tissue volume after delineating PR cells from non-PR cells. Elav revealed a rosette-like pattern of DAPI stained nuclei in PR cells that were used for RNA counts (Supp. Fig. 2A). As also seen in the images (Fig. 1), we see that *spitz* mRNA is mostly expressed in the photoreceptor cells (Elav-positive cells) whereas *argos* mRNA is highly expressed in the non-photoreceptor cells, Elav-negative cells (Fig. 1C). The directionality in signaling can also be visualized by dpERK staining, a classical marker for high EGFR signaling, in the non-PR cells (Supp. Fig. 2B). mRNA counts quantified along a line from the morphogenetic furrow to the posterior end of the eye imaginal disc do not show a large variation along tissue length (Supp. Fig. 2C and Fig. 1D) indicating that the expression patterns are stable in time.

**Figure 1.**
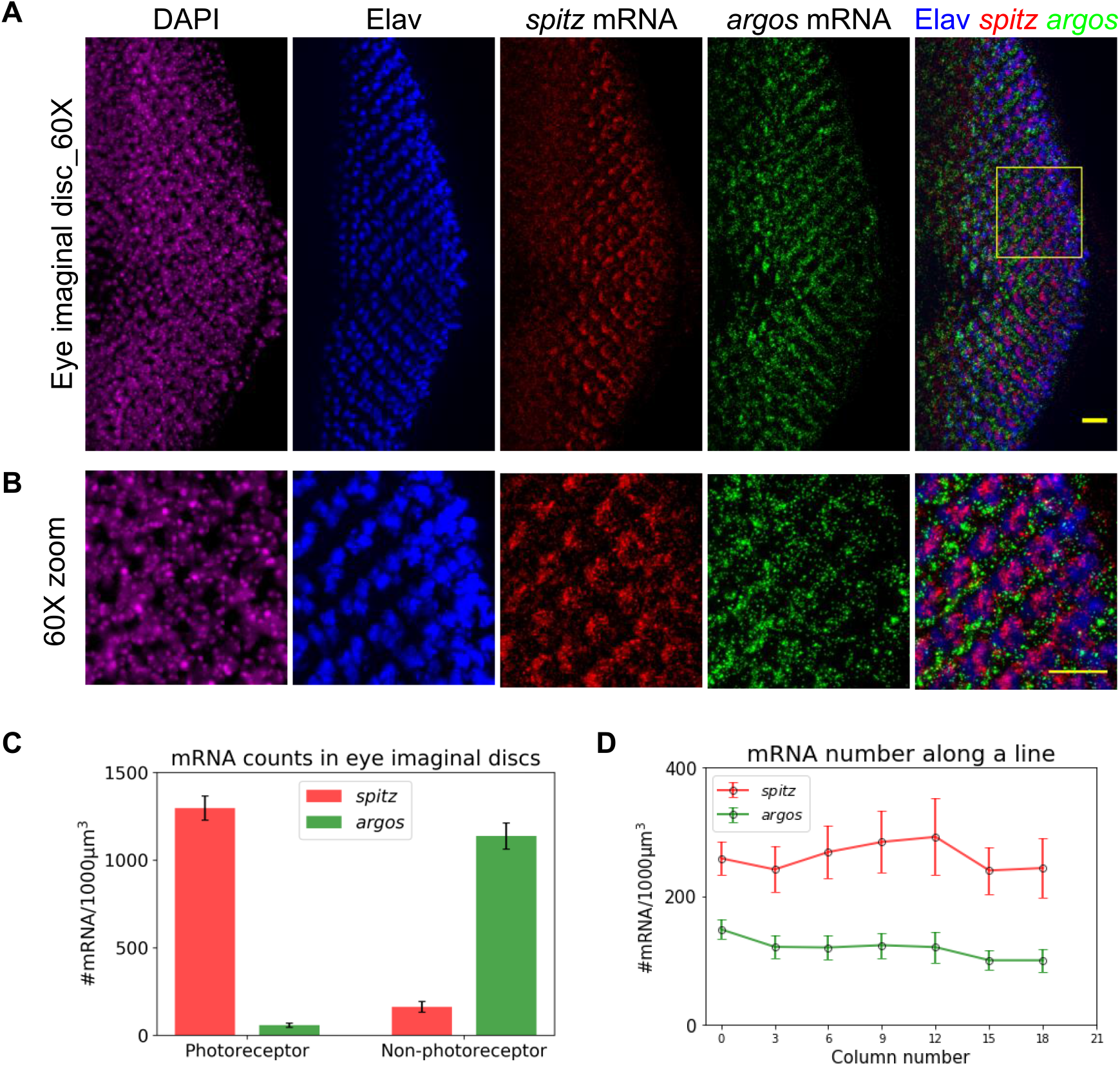
EGFR signaling is directional in the eye imaginal discs. (A) 60X images of 3^rd^ instar eye imaginal discs stained with DAPI, Elav (Immunofluorescence; pan-neuronal marker), *spitz* mRNA and *argos* mRNA (single molecule RNA FISH). *spitz* mRNA is highly expressed only in the Elav-positive photoreceptor cells, whereas *argos* mRNA is highly expressed in the neighbouring non-neuronal cells. To clearly see the exclusive expression of *spitz* and *argos*, the image is zoomed into the yellow box and presented in (B). Scale bar is 10μm in (A) and 5 μm (B). These are z-projected images and hence individual transcripts get merged. A single z-slice is shown in Supp. Fig. 1. (C) The individual transcript molecules are counted in 3D using the StarSearch software (https://www.seas.upenn.edu/~rajlab/StarSearch/launch.html). The respective counts are represented per 1000 μm^3^ of tissue volume as individual cells are hard to segment in this dense tissue. Photoreceptor (PR) and non-photoreceptor (non-PR) cells are identified by nuclei position corresponding to Elav as shown in Supp. Fig. 2A. (N=9 tissues) (D) The mRNA counts are also represented in each photoreceptor column along a line from morphogenetic furrow (column 0) to the posterior end of the eye imaginal disc, as indicated in Supp. Fig. 2C. (N=8 tissues), irrespective of cell type (PR or non-PR) - overall mRNA counts are relatively constant along the line i.e. with time after PR differentiation. The error bars in (C) and (D) are standard errors of mean.

### Effects of modulating transcript numbers on eye phenotype

Knocking down components of the DER pathway during development in the eye imaginal disc is known to affect the adult eye phenotype. It has been suggested that the ratio of Spitz to Argos may be crucial for proper eye patterning. Halving Spitz in the background of hypomorphic Argos reverted the rough eye caused by the latter whereas in the background of Argos overexpression, it enhanced the rough phenotype (Schweitzer et al., 1995b). We used the UAS-Gal4 system (Brand and Perrimon, 1993; Fischer et al., 1988) to knockdown *argos* and *spitz*. GMR-Gal4 (Li et al., 2012) was used to drive dsRNA cassettes in all the cells posterior to the morphogenetic furrow and Elav-Gal4 (Yao and White, 1994) was used to drive the same cassettes in the photoreceptor (neuronal) cells. smFISH showed the decrease in respective mRNA transcripts compared to wildtype CantonS eye discs. *argos* expression was decreased even when *spitz* was knocked down being downstream in the cascade triggered by Spitz (Fig. 2A; two rows from above). Expectedly when either of the genes were knocked down using a GMR-Gal4 driver in all cells, the flies showed a fully penetrant rough eye phenotype; but to our surprise while the Elav-Gal4 driver did show knockdown of the *spitz*, there was absolutely no discernible phenotype in the adult eye. The SEM images of adult fly eyes had normal ommatidial patterns when *spitz* was knocked down using a Elav-Gal4 driver (Fig. 2A; lower panel). As expected, eye discs did not show any change in transcript numbers when *argos* dsRNA was driven by Elav-Gal4, nor was there any defect in the adult eye. *argos* and *spitz* mRNA was quantified in all the crosses along with CantonS and the numbers reflected the respective gene knockdowns (Fig. 2B and 2C). The quantified gene expression could not explain the absence of effect on adult eye phenotype in the *spitz* knockdown driven by Elav-Gal4. As relative dosage of EGFR pathway is known to be important for photoreceptor differentiation (Schweitzer et al., 1995b; Taguchi et al., 2000), we wondered if the ratio of expression of *spitz* to *argos* may be the critical determinant of the final eye phenotype instead of absolute levels of expression. This does assume that mRNA level expression translates directly to levels of final activated protein products, but given that Spitz and Argos act in a 1:1 stoichiometry, we thought it would be a reasonable idea to test. We analyzed the *spitz*-to-*argos* expression ratio in the above crosses in the eye discs (with no distinction of PR and non-PR). Remarkably, the *spitz*-to-*argos* ratio in the progeny of Elav-Gal4 with *spitz* dsRNA was similar to wildtype CantonS flies, while it was either significantly higher or lower in other crosses (Fig. 2D). Thus, the gene expression ratio of the activating ligand and the negative feedback molecule might dictate the final availability of free active ligand for pathway activation rather than their absolute expression levels.

**Figure 2.**
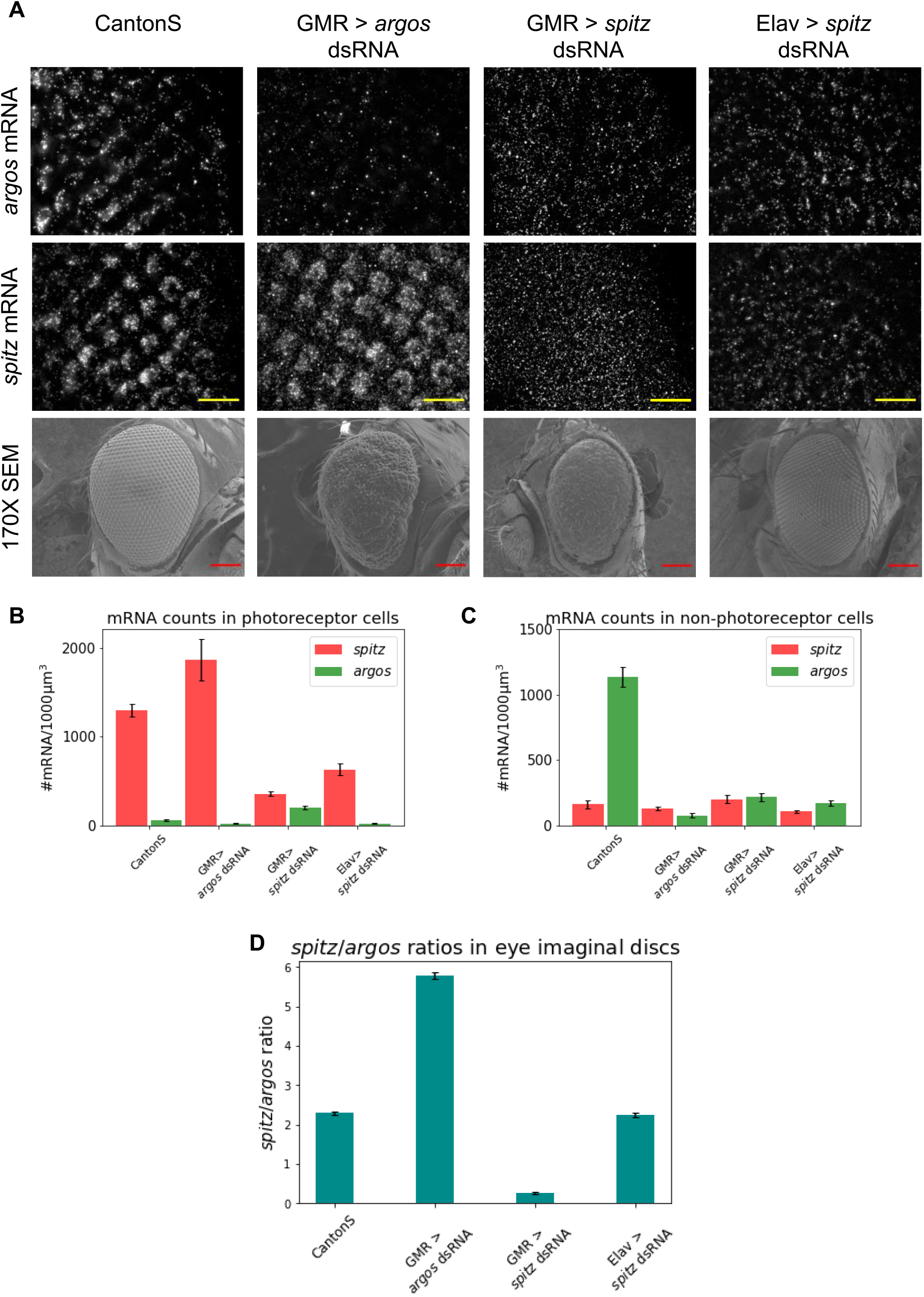
*spitz*-to-*argos* ratio is important for proper ommatidial patterning. (A) smFISH was performed in eye imaginal discs from represented crosses for *spitz* and *argos* mRNA in the two rows. In the third row, adult eye phenotypes are shown. The Elav > UAS *spitz* dsRNA progeny shows perfectly arranged ommatidia like the wildtype CantonS, while the other two crosses show rough eye phenotypes. Scale bar for widefield fluorescence images is 10μm and SEM images is 100μm. (B) and (C) mRNA numbers for *spitz* and *argos* were counted in photoreceptors and non-photoreceptors in respective crosses. *spitz* and *argos* mRNA show variation in number as expected from the respective crosses. (D) *spitz*-to-*argos* mRNA ratio in eye field irrespective of cell type (PR or non-PR) was calculated as dosage was known to be important for ommatidial pattern formation (N=9 tissues in all the crosses). Most remarkably, knocking down *spitz* in PR cells with a Elav driver, knocks down *argos* numbers too, and overall the ratio is unchanged. When a GMR driver is used to knockdown either *argos* or *spitz* in the full field, the ratio is higher or lower than wildtype. Error bars in (B), (C) and (D) are standard errors of mean.

### Buffered *spitz*-to-*argos* ratio is important for proper ommatidial patterning

To test the hypothesis that the ratio of expression of *spitz*-to-*argos* is critical for determining the final eye phenotype we attempted to use other cassettes to modulate their levels and also to tune their expression in a systematic manner. EGFR^CA^ is a constitutive active form of DER, and hence the downstream targets will be expressed throughout the development. We used a GMR-Gal4 driver to express this constitutive DER. As GMR expresses in all the cells posterior to the morphogenetic furrow (Freeman, 1996), the patterned expression of *argos* is lost (Fig. 3A). *spitz*-to-*argos* expression ratio was quantified for the EGFR^CA^ progeny, the balancer control (non-EGFR^CA^) and CantonS larvae. The balancer control had *spitz*-to-*argos* ratio levels similar to the CantonS around 2-2.2, whereas the EGFR^CA^ larvae (as determined from tubby marker) had decreased ratio around 0.5 given high expression of *argos* (Fig. 3B). This ratio change was reflected in the adult eye SEM images (Fig. 3C). EGFR^CA^ had dramatically smaller and rough eye. To investigate if patterning defects in the adult eye vary continuously with the *spitz*-to-*argos* ratio or show up beyond a specific threshold indicative of developmental buffering, we tuned the expression of *spitz* and *argos*. We generated a GMR-Gal4 line with a temperature sensitive Gal80 (Lee and Luo, 1999; McGuire et al., 2003) to drive different cassettes affecting DER pathway components. Gal80^ts^ binds to Gal4 and inhibits transcriptional activation at 18°C but turns inactive at 29°C. EGFR^CA^ was crossed with this new driver line and shifted to 29°C for different time points. The larvae were processed for smFISH and the absolute gene expression for *spitz* and *argos* (Fig. 3D) and the *spitz*-to-*argos* ratios (Fig. 3E) were quantified. The corresponding SEM images of the adult eyes were compared. The adult eye phenotype started to show patterning defects when the *spitz*-to-*argos* gene expression ratios neared 1 upon 40 minutes of heat shock at 29°C (Fig. 3F).

**Figure 3.**
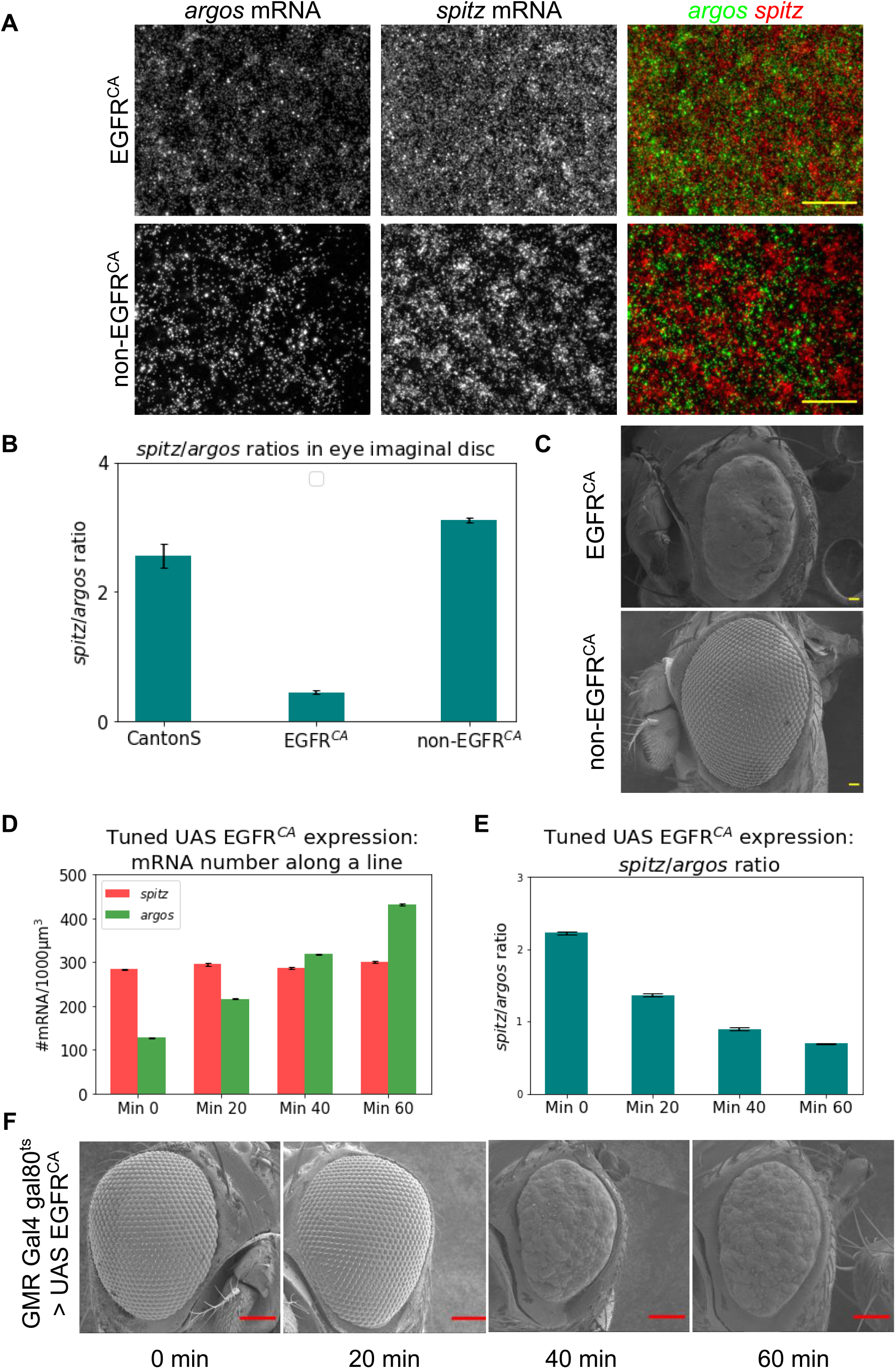
Overexpression of EGFR completely disrupts ommatidial pattern from the adult eye. (A) Eye imaginal discs of GMR > UAS EGFR^CA^/TM6 progeny stained for *spitz* and *argos* mRNA. From the same cross, EGFR^CA^ and non-EGFR^CA^ larvae were distinguished using the tubby phenotype. Scale bar for widefield images is 10μm. Exclusive expression of *spitz* and *argos* is lost in EGFR^CA^ which leads to change in the *spitz*-to-*argos* ratio represented in (B) [CantonS N=9, EGFR^CA^ N=10, non-EGFR^CA^ N=9]. (C) 170X SEM images of GMR > UAS EGFR^CA^/TM6 progeny. The ommatidial pattern is lost when EGFR is constitutively active and the size of the eye is also reduced. Heatshock was administered for 6 hrs to EGFR^CA^ and non-EGFR^CA^ larvae for maximal activation of the cassette. Scale bar is 10μm. (D) 170X SEM images of GMR > UAS EGFR^CA^ adult flies corresponding to different time points of heat shock at 29°C. N=8 for all time points. (E) *spitz*-to-*argos* ratios in eye discs corresponding to different time points of heat shock at 29°C. Quantification of absolute mRNA numbers and ratios represented in all the above plots were calculated in the eye field irrespective of the cell type. Error bars in all the plots are standard errors of mean. (F) Adult eyes corresponding to different time points of heat shock at 29°C. Scale bar is 100μm. No rough-eye phenotype is seen for 20 min of heatshock, though a difference in the *spitz*-to-*argos* ratio is already seen at this point. Beyond this the eyes are fully rough, and the phenotype is fully penetrant. This is indicative of developmental buffering.

### Threshold switch of *spitz*-to-*argos* ratio regulates phenotype

To recapitulate the buffer range with another cassette, we crossed UAS Argos to the GMR-Gal4 Gal80^ts^ driver line to overexpress a*rgos*. Absolute gene expression (Fig. 4A) and ratio (Fig. 4B) were quantified. The corresponding SEM images (Fig. 4C) were observed for patterning defects. The adult eyes showed patterning defects when the *spitz*-to-*argos* ratio neared 1. This confirmed that the adult eye phenotype is indeed contingent on the *spitz*-to-*argos* ratio. The phenotype did not show any defects in the adult eye even though the *spitz*-to-*argos* ratio dropped from 2.2 in the no-heat shock control till it reaches 1. We did not capture any defects in the ommatidial patterning in the flies with intermediate ratio. This suggests that patterning defects show up only when the *spitz*-to-*argos* ratio crosses a certain threshold and the intermediate ratios are buffered in the development. Having investigated the effects of decreasing the *spitz*-to-*argos* ratio, we wondered how far the buffering will persist when we increased it from wildtype levels. For this we used an EGFR^DN^ (dominant negative allele of DER) cassette with the GMR-Gal4 Gal80^ts^ driver, where ratios would be driven to values greater than wildtype. *spitz* and *argos* mRNA transcripts were quantified (Fig. 4D) along with their ratio (Fig. 4E) at different time points of heat shock at 29°C. The respective SEM images (Fig. 4F) showed ommatidial defects when the ratio approached 3. The buffer range of *spitz*-to-*argos* for contributing towards proper ommatidial patterning might lie between 1-3 while the wildtype ratio lies around 2-2.2. The tight range for wildtype and balancer controls is perhaps indicative of the importance of relative Spitz and Argos doses, but the system is developmentally buffered to thresholds beyond this wildtype range. We never saw mixed phenotypes, and the flies always showed either largely proper ommatidial arrangements, or fully penetrant rough-eye phenotypes.

**Figure 4.**
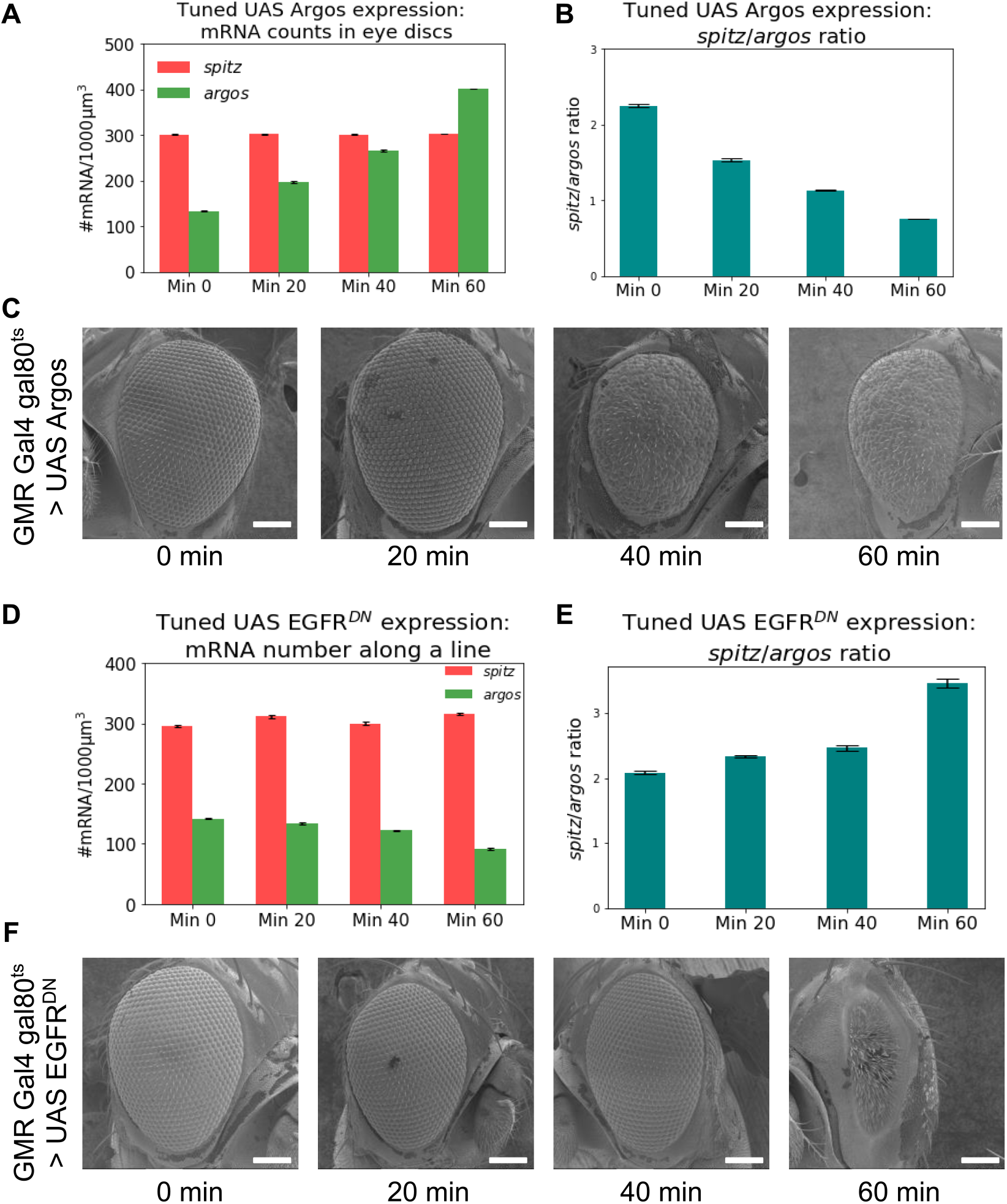
Binary switch in *spitz*-to-*argos* threshold range for proper patterning of ommatidia. (A) Absolute *spitz* and *argos* mRNA numbers from GMR-Gal4 Gal80^ts^ > UAS Argos progeny after different time points of heat shock at 29°C. (B) *spitz*-to-*argos* ratios from eye discs of GMR-Gal4 Gal80^ts^ > UAS Argos progeny after different time points of heat shock at 29°C. N=9 tissues for all time points. (C) 170X SEM images of adult eyes of GMR-Gal4 gal80^ts^ > UAS Argos progeny at different time points of heat shock at 29°C. (D) Absolute *spitz* and *argos* mRNA numbers from GMR-Gal4 Gal80^ts^ > UAS EGFR^DN^ progeny after different time points of heat shock at 29°C. (E) *spitz*-to-*argos* ratios from eye discs of GMR-Gal4 Gal80^ts^ > UAS EGFR^DN^ progeny after different time points of heat shock at 29°C. N=8 tissues for all time points. Quantification of absolute mRNA numbers and ratios represented in all the above plots were calculated in the eye field irrespective of the cell type. Error bars for all the plots are standard errors of mean. (F) 170X SEM images of adult eyes of GMR-Gal4 gal80^ts^ > UAS EGFR^DN^ progeny at different time points of heat shock at 29°C. Scale bar for all SEM images is 100μm. In both cases, the final adult phenotype is robust to certain variation in the ratio threshold, and breaks down beyond resulting in fully penetrant rough-eye phenotypes. It should be noted that qualitatively the roughness phenotype looks different between UAS Argos and UAS EGFR^DN^.

## Discussion

Developmental pathways have evolved mechanisms to monitor positional information in order to generate reproducible organismal patterns. These pathways are robust and insensitive to small changes in individual processes involved. Spatial differentiation, where a population of cells undergo deterministic molecular differentiation, brings about spatial patterns (Wolpert, 1969). Redundancy of mechanism and negative feedback are two ways in which reliability in pattern formation is brought about. Lateral inhibition by diffusible molecules is another mechanism that can be used to generate patterns. For systems which do not depend on developmental history, environmental makeup determines their molecular differentiation contributing towards generating a pattern.

In this paper, we propose a mechanism where relative expression levels of principal EGFR ligand, Spitz and the feedback molecule, Argos determines the extent of DER activation which is crucial for periodic ommatidial pattern. We show that absolute gene expression may not be critical but the balance between gene networks on the whole may contribute towards proper pattern formation. The eye discs expressing EGFR^CA^ construct under GMR-Gal4 Gal80^ts^ with *spitz*-to-*argos* ratio near 1, showed discontinuous S-phase band after the morphogenetic furrow indicating a lower population of cells entering the second mitotic wave (SMW) compared to the no-heat shock control (Supp. Fig. 3A). Fewer cells entering the SMW leaves the tissue field with fewer uncommitted cells to make cell fate decisions. This can affect pattern formation to a great extent. This could also explain fewer number of bristle cells in the rough adult eyes (Fig. 3 and Supp. Fig. 3B) as bristles cell fate is assigned from cells arising from the second mitotic wave (De Nooij and Hariharan, 1995). It should also be noted that the rough eye phenotype for EGFR^DN^ is rather different from EGFR^CA^ and shows a profusion of bristles (compare 60 min time-points of Fig. 4F and Fig. 3F). This mechanism of relative expression determining phenotype supports older work on the importance of *spitz*-to-*argos* dose as a critical determinant of eye patterning (Schweitzer et al., 1995b; Taguchi et al., 2000), and leads to new knowledge about differential expression of these genes in the early eye field. This work also addresses the sensitivity of the system to the heterogeneity in the expression levels of gene networks and makes developmental programs robust. It has to be noted of course that signaling via EGFR is not the only pathway determining the ommatidial pattern in the eye. For example, Notch is known to play an important role in initiation of neural development and also for ommatidial rotation (Baker and Zitron, 1995; Baonza and Freeman, 2001). Buffered regulation of genes in different developmental pathways that crosstalk can decrease sensitivity to variations in a gene network and can help explain other reproducible and stereotypical patterns generated throughout the development.

## Materials and Methods

### Drosophila stocks and crosses

All fly strains were grown on standard cornmeal agar at 25°C. CantonS line was used as the wildtype fly. The cassettes used for modulating the DER pathway were UAS argos dsRNA (GD47180), UAS spitz dsRNA (BL34645), EGFR^CA^/TM6, homozygous UAS Argos and EGFR^DN^ (2,3). Tissue specific GMR and Elav drivers were used to express the above constructs. Gal80^ts^/FM7; sp/CyO was used to generate GMR-Gal4 with a temperature sensitive gal80. All crosses with Gal80^ts^ line were grown at 18°C, and heat shock was given at 29°C unless otherwise mentioned.

### Tissue preparation and smFISH

3rd instar larvae were collected in nuclease-free 1X PBS (Ambion, AM9624) and washed once with the same. The larvae were flipped and transferred to a microcentrifuge tube containing 4% Paraformaldehyde (PFA, Sigma, P6148) in 1X PBS for 25 minutes at room temperature. The fixative was aspirated and washed twice with 1X PBS. The tissue was permeabilized using 0.3% Triton-X 100 (Sigma, T8787) in 1X PBS for 45 minutes at room temperature. The permeabilizing agent was aspirated and the tissue washed twice with 1X PBS. The tissues were kept in 70% ethanol at 4°C overnight. The tissues were washed twice with a wash buffer (20% Formamide (Ambion, 9342) and 2X SSC (Ambion, AM9763) in nuclease-free water) for 30 minutes each at 37°C. The wash buffer was aspirated and the tissues were incubated with a hybridization mix overnight at 37°C. The probe sequences for *spitz* and *argos* is given in the previous paper (Pasnuri et al., 2018). The mix was removed and tissues were washed twice with wash buffer at 37°C. DAPI (Invitrogen, D1306; 2μg/ml) in wash buffer was added to the tissues and incubated for 30 minutes at 37°C. The tissues were then washed with 2X SSC twice for 5 minutes. The eye imaginal discs were dissected in 2X SSC and mounted on a clean slide with a drop of Vectashield (Vecta Labs). A cartoon depicting singly labeled 20-mer oligonucleotide probes binding to mRNA target is shown in Supp. Fig. 1A. StarSearch software (https://www.seas.upenn.edu/~rajlab/StarSearch/launch.html) was used to count transcripts in 3D as done before (Pasnuri et al., 2018). The absolute counts and ratios were calculated in regions of interest on the eye field and then averaged.

### Antibody staining

The tissues were fixed and permeabilized as described earlier. The tissues were then washed twice with 1X PBS. Blocking solution (5% BSA in 1X PBS) was added to the tissues and incubated for 1 h at room temperature on a rotating mixer. Primary antibody (1:1000 Elav [DSHB, 7E8A10], 1:500 Yan [DSHB, 8B12H9], 1:500 Rough [DSHB, 62C2A8], 1:1000 pH3 [CST, 9701S]) diluted in the blocking solution (5% BSA [Sigma, A2153] in 1X PBS) was added after 1hour and incubated overnight at 4°C. 1:1000 secondary antibody (Goat anti-Rat 488 [Invitrogen, A11006], Goat anti-Rabbit 594 [Invitrogen, A11037], Goat anti-Mouse [Invitrogen, A11029]) diluted in blocking solution was added to the tissues and incubated for 3 hours at room temperature. Tissues were then washed twice with 1X PBS for 15 minutes each on a rotating mixer. DAPI in 1X PBS was added and incubated for 30 minutes. The tissues were finally washed twice with 1X PBS for 5 minutes each.

For simultaneous smFISH-IF, the primary antibody is added into the hybridization mix and incubated overnight at 37°C. The secondary antibody was diluted to 1:1000 in nuclease-free blocking solution and incubated with tissues for 3 hours at room temperature on a rotating mixer.

### Image acquisition

The tissues were imaged on an Olympus BX63 upright widefield fluorescence microscope with a Retiga 6000 (Qimaging) CCD monochrome camera. The slides were kept at 4°C for 1 hour before imaging. The images were acquired using a 60X, 1.42 N.A. oil immersion objective with a z-step size of 0.3μm. Narrow band-pass filters (ChromaTechnology −49309 ET- Orange#2 FISH for Quasar 570 labelled probes; 49310 ET- Red#2 FISH for CAL fluor 610 labelled probes) were used to spectrally separate single transcripts imaged in two colors.

### SEM imaging of adult eyes

Adult fly heads were dissected in 1XPBS. PBS was aspirated and 4%PFA was added for 25 minutes at room temperature. Fly heads were then washed twice with 1X PBS for 5 minutes each. Conducting carbon adhesive tape was spread on the SEM specimen stub. The fly heads were aligned on the carbon tape and were air-dried. The images were captured using a lower electron detector in a JEOL JSM7200F microscope at 170X magnification.

## Funding and acknowledgements

We thank Dr. Rohit Joshi and Dr. Rakesh Mishra for the gift of fly lines. We also thank Dr. Manish Jaiswal for sharing fly resources and valuable discussions. We also thank Shravani Anagandula for helping us with SEM imaging. This project was supported by intramural funds at TIFR Hyderabad from the Department of Atomic Energy, Government of India (Project Identification No. RTI 4007).

## Supplementary data

**Supplementary Figure 1:**
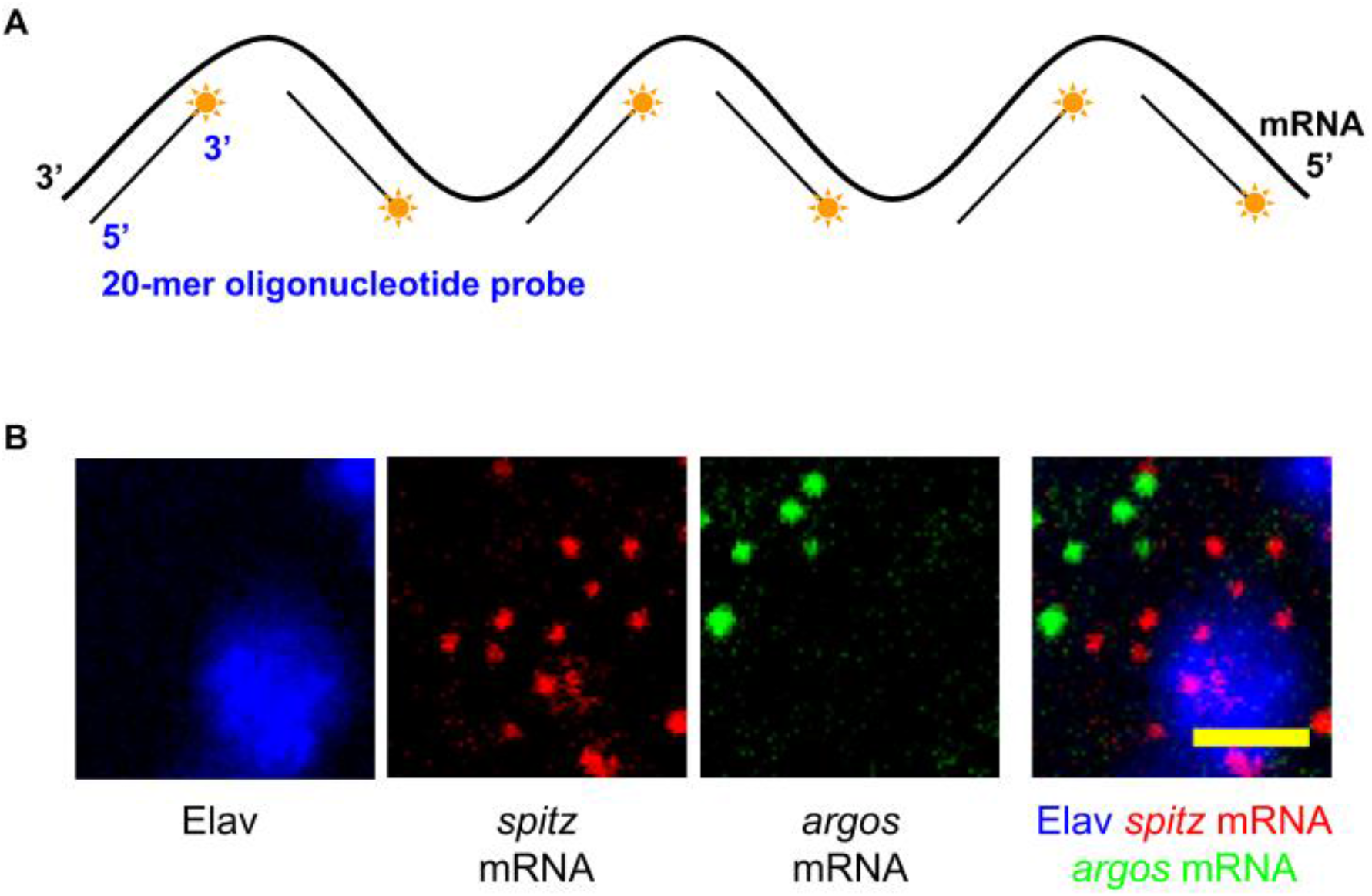
(A) Cartoon showing singly-labeled 20-mer probes binding to mRNA in single molecule RNA FISH protocol. Multiple 20 nt long oligos each carrying a 3′ fluorophore is used to decorate the mRNA of interest following previous protocols. (B) Zoomed images of a single z-slice showing single transcripts of *spitz* mRNA and *argos* mRNA. Scale bar is 2μm.

**Supplementary Figure 2:**
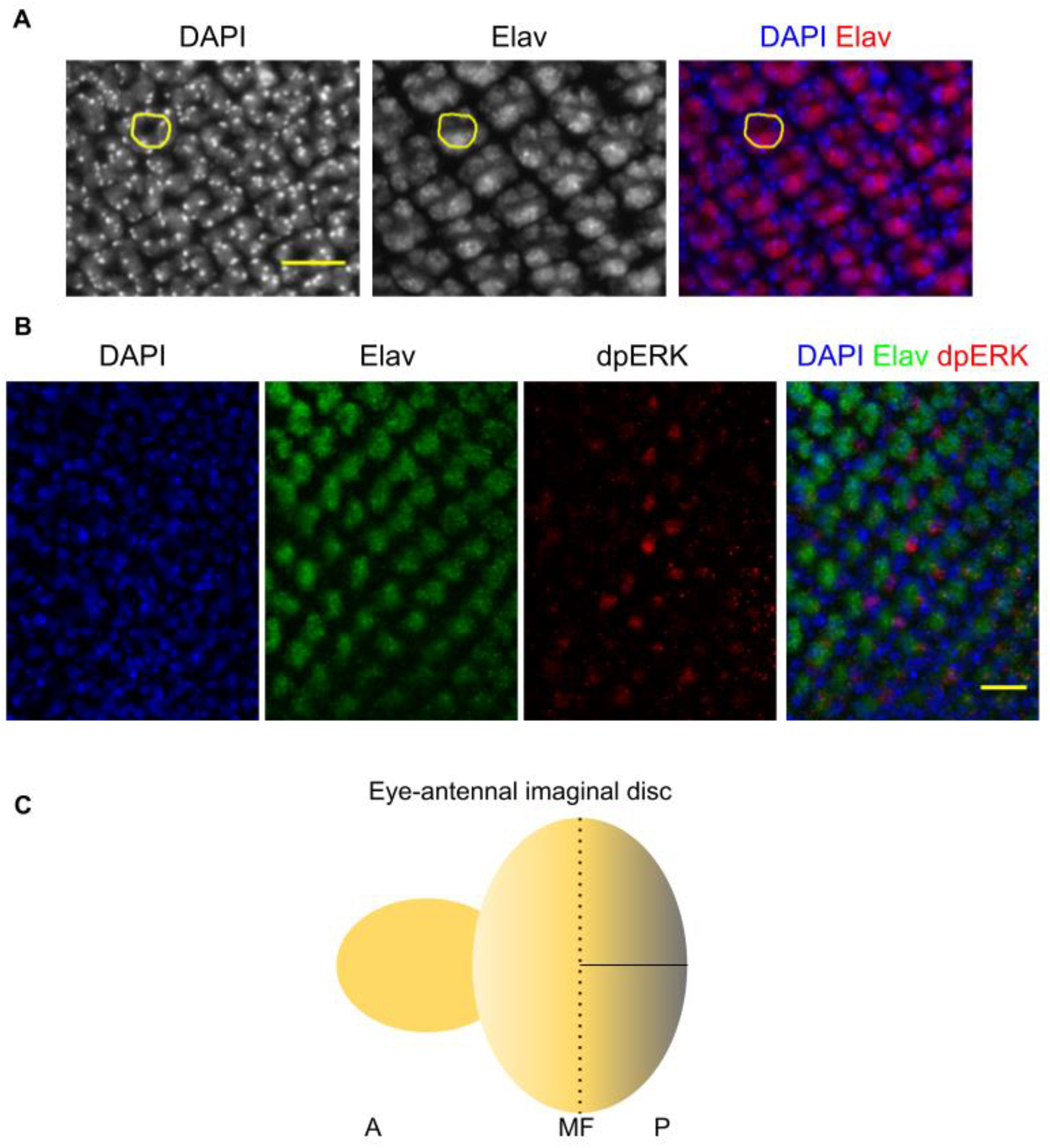
(A) Rosette like PR nuclei are stained by a pan-neuronal marker, Elav. (B) The Elav-positive cells and cells stained with dp-ERK are exclusive, again saying that EGFR signaling is activated in the neighbouring cells to the ligand source of PR cells. (C) Cartoon representing the line along which *spitz* and *argos* mRNA counts were analysed (A:

**Supplementary Figure 3:**
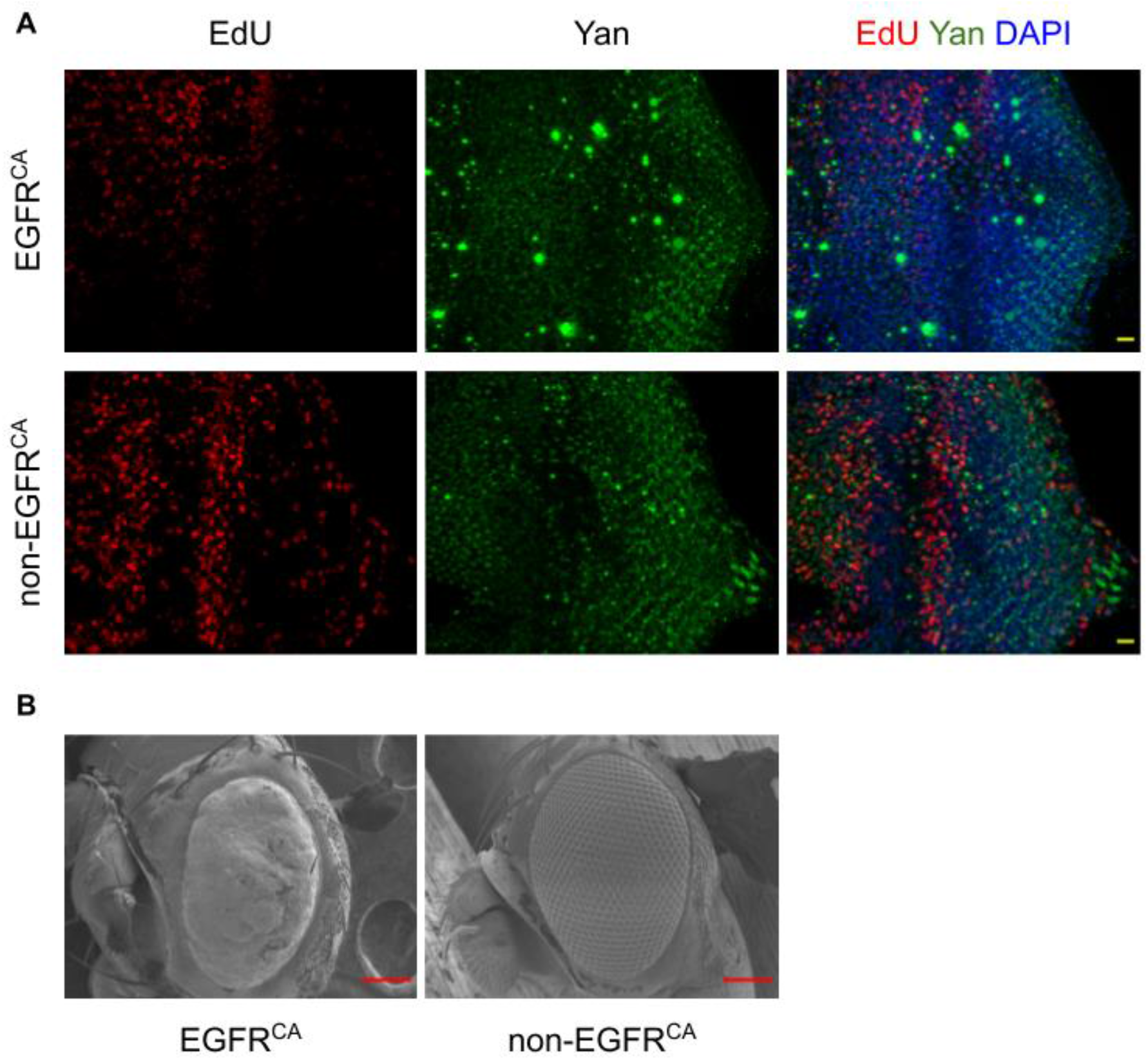
(A) EdU staining which mark the cells re-entering the cell cycle in the second mitotic wave is discontinuous in the EGFR^CA^ eye discs but form a continuous band behind the morphogenetic furrow in non-EGFR^CA^ eye discs, similar to wildtype discs. Scale bar is 10μm. (B) In this experiment the flies were shifted from room temperature (instead of 18°C) to 29°C for 6 hrs. But the adult eye phenotypes are same as that seen in Fig. 3C. SEM images of adult eyes are presented here. Scale bar is 100μm.

